# Punishment resistance for cocaine is associated with inflexible habits in rats

**DOI:** 10.1101/2023.06.08.544242

**Authors:** Bradley O. Jones, Morgan S. Paladino, Adelis M. Cruz, Haley F. Spencer, Payton L. Kahanek, Lauren N. Scarborough, Sandra F. Georges, Rachel J. Smith

**Affiliations:** Institute for Neuroscience, Texas A&M University, College Station, TX, USA; Department of Psychological and Brain Sciences, Texas A&M University, College Station, TX, USA

## Abstract

Addiction is characterized by continued drug use despite negative consequences. In an animal model, a subset of rats continues to self-administer cocaine despite footshock consequences, showing punishment resistance. We sought to test the hypothesis that punishment resistance arises from failure to exert goal-directed control over habitual cocaine seeking. While habits are not inherently permanent or maladaptive, continued use of habits under conditions that should encourage goal-directed control makes them maladaptive and inflexible. We trained male and female Sprague Dawley rats on a seeking-taking chained schedule of cocaine self-administration (2 h/day). We then exposed them to 4 days of punishment testing, in which footshock (0.4 mA, 0.3 s) was delivered randomly on one-third of trials, immediately following completion of seeking and prior to extension of the taking lever. Before and after punishment testing (4 days pre-punishment and ≥4 days post-punishment), we assessed whether cocaine seeking was goal-directed or habitual using outcome devaluation via cocaine satiety. We found that punishment resistance was associated with continued use of habits, whereas punishment sensitivity was associated with increased goal-directed control. Although punishment resistance was not predicted by habitual responding pre-punishment, it was associated with habitual responding post-punishment. In parallel studies of food self-administration, we similarly observed that punishment resistance was associated with habitual responding post-punishment but not pre-punishment. These findings indicate that punishment resistance is related to habits that have become inflexible and persist under conditions that should encourage a transition to goal-directed behavior.

## Introduction

Addiction is characterized by compulsive drug seeking and continued drug use despite negative consequences. In an animal model of compulsive drug use, a subset of rats continues to self-administer cocaine despite footshock consequences, indicating punishment resistance (Belin *et al*, 2008; Deroche-Gamonet *et al*, 2004; Pelloux *et al*, 2007; Vanderschuren and Everitt, 2004). Compulsive drug use has been theorized to stem from a loss of control over habitual behavior, making habits maladaptive and inflexible (Belin *et al*, 2013; Brown *et al*, 2022; Everitt and Robbins, 2005, 2016; Everitt, 2014; Ostlund and Balleine, 2008; Smith and Laiks, 2018). Although habits are considered automatic and insensitive to changes in outcome value, they are not necessarily permanent or insensitive to consequences. Rather, habitual behavior is typically flexible in that it is overridden by goal-directed control under conditions of punishment or changes in context (Bouton, 2021; Ostlund and Balleine, 2008). In contrast, habitual responding that persists despite conditions that should encourage goal-directed control may indicate that habits have become maladaptive and inflexible. Here we sought to directly assess the relationship between habitual cocaine seeking and punishment resistance in rats.

The role of cocaine-seeking habits in the development of punishment resistance has been unclear, partially due to limited methods for assessing habitual responding for intravenous (IV) cocaine. We recently developed a procedure to discriminate goal-directed and habitual responding in rats self-administering IV cocaine using outcome devaluation via satiety (Jones *et al*, 2022). Goal-directed behavior is performed in direct pursuit of the outcome, and therefore sensitive to outcome devaluation, whereas habitual behavior is automatically elicited by conditioned stimuli and insensitive to outcome devaluation (Balleine and Dickinson, 1998; Dickinson, 1985; Yin and Knowlton, 2006). Using this novel outcome devaluation procedure, we found that bilateral lesions of dorsolateral striatum (DLS) or dorsomedial striatum (DMS) caused goal-directed or habitual cocaine responding, respectively, similar to previous work with food rewards (Balleine and O’Doherty, 2010; Corbit and Janak, 2010; Gremel and Costa, 2013; Jones *et al*, 2022; McNamee *et al*, 2015; Yin and Knowlton, 2006; Yin *et al*, 2004, 2006). An advantage of this procedure is that it elicits devaluation temporarily without the need for additional training, easily allowing repeated testing at different time points (e.g., before and after footshock punishment testing).

While habitual responding develops in the majority of rats after extended training on cocaine self-administration (Leong *et al*, 2016; Zapata *et al*, 2010), punishment resistance develops in only a subset of rats (Deroche-Gamonet *et al*, 2004; Jonkman *et al*, 2012a; Pelloux *et al*, 2007). DLS is necessary for habitual responding for cocaine and is progressively recruited over extended cocaine training (Bender and Torregrossa, 2023; Jones *et al*, 2022; Murray *et al*, 2012; Porrino *et al*, 2004a, 2004b; Willuhn *et al*, 2012; Zapata *et al*, 2010). DLS may also play a role in punishment resistance for cocaine, considering that DLS inactivation increased sensitivity to footshock punishment (Jonkman *et al*, 2012b). Similar parallels between habits and punishment resistance have been observed for alcohol. Extended alcohol exposure increased habitual responding and DLS control of self-administration, as well as punishment resistance despite footshock (Corbit *et al*, 2012, 2014; Giuliano *et al*, 2019, 2021; Lopez and Becker, 2014; Radke *et al*, 2017). Animals whose alcohol seeking had become habitual and DLS-dependent after extended training showed continued alcohol seeking despite footshock, supporting a role for habits in punishment resistance (Giuliano *et al*, 2019, 2021). In contrast, while extended training with food rewards leads to increased use of habits, it has not been shown to increase punishment resistance (Adams, 1982; Dickinson *et al*, 1995; Limpens *et al*, 2014; Pelloux *et al*, 2007, 2015; Radke *et al*, 2017; Vanderschuren and Everitt, 2004). In summary, extended training with cocaine or alcohol results in habitual responding in the majority of animals, as well as punishment resistance in a subset of animals, and these two processes may be linked in addiction. Alternative theories posit that addiction is driven by excessive goal-directed choice and/or overvaluation of drug reward, and that habits are not necessary (Hogarth, 2020; Singer *et al*, 2018).

To investigate the relationship between habitual cocaine seeking and punishment resistance, we trained male and female rats to self-administer IV cocaine on a seeking-taking chained schedule of reinforcement, originally developed by Olmstead et al (2000, 2001) and used extensively to study punishment resistance (Chen *et al*, 2013; Datta *et al*, 2018a, 2018b; Giuliano *et al*, 2018, 2019, 2021; Jonkman *et al*, 2012a; Limpens *et al*, 2014, 2015; Pelloux *et al*, 2015, 2007; Vanderschuren and Everitt, 2004; Zhou *et al*, 2019). We then exposed rats to 4 days of punishment testing, and used outcome devaluation via cocaine satiety to assess whether responding was goal-directed or habitual 4 days pre-punishment and at least 4 days post-punishment. We found that punishment resistance for cocaine was associated with habitual responding post-punishment but not pre-punishment. In parallel experiments in which rats were trained to self-administer food on the same schedule, we also found that punishment resistance was associated with habitual responding post-punishment but not pre-punishment. These data indicate that punishment resistance is associated with inflexible habits, whereas punishment sensitivity is associated with increased goal-directed control.

## Methods

### Animals

Male and female Sprague Dawley rats (initial weight 225-250 g; Charles River, Raleigh, NC, USA) were single-housed in a temperature- and humidity-controlled facility at Texas A&M University accredited by AAALAC. Rats were housed under a reversed 12-h light/dark cycle (lights off at 6 a.m.), with food and water access ad libitum, except when noted below. All experiments were approved by the IACUC at Texas A&M and conducted according to specifications of the NIH as outlined in the Guide for the Care and Use of Laboratory Animals.

### Surgery

For cocaine self-administration studies, rats were anesthetized via isoflurane (induction 5%, maintenance 1-3%), given a nonsteroidal anti-inflammatory analgesic (ketoprofen, 2 mg/kg, s.c.), and implanted with chronic indwelling IV jugular catheters, as previously described (Smith *et al*, 2009). Beginning three days after surgery, catheters were flushed once daily with 0.1 ml of cefazolin (100 mg/ml) and 0.1 ml heparin (500 U/ml). Self-administration sessions began after at least 5 days of recovery from surgery.

### Cocaine self-administration

Rats were trained to self-administer IV cocaine (0.5 mg/kg per infusion) on a seeking-taking chained schedule of reinforcement, in which completion of a random ratio (RR20) or random interval (RI60) schedule on the seeking lever gave access to the taking lever during daily 2-h sessions. Infusions of cocaine (pump speed of 70 μg/s) were paired with 5-sec tone and light cues (78 dB, 2900 Hz; white stimulus light above the active lever). Operant conditioning chambers were housed in sound-attenuating cubicles and controlled via MED-PC IV (Med-Associates, St. Albans, VT). Cocaine HCl was obtained as a gift through the NIDA Drug Supply Program.

To train animals, self-administration began with fixed ratio (FR) 1 reinforcement, with only the taking lever available (criterion of 5 sessions ≥20 infusions). Rats were food-restricted (85-90% of free-feeding weight) at the start of the experiment to increase general motivation, and were placed back onto free feeding once they had at least 2 consecutive sessions where they earned ≥20 infusions. Training then progressed to a chained seeking-taking schedule with FR1 (seeking) - FR1 (taking) reinforcement (criterion of 2 days ≥15 infusions), during which completion of the seeking link of the chain led to retraction of the seeking lever and extension of the taking lever; completion of the taking link of the chain delivered cocaine and led to retraction of the taking lever and the start of the next trial. During the seeking link of the chain, a stimulus light (S+) was presented above the seeking lever and signaled availability. At the next stage, rats were given a 4-min time out between trials, such that completion of the taking link of the chain led to retraction of the taking lever and extension of the seeking lever, but with no S+ and no programmed consequence for responding (criterion of 2 days ≥10 infusions). Training then progressed to RR or RI seeking schedules, and the taking lever was available for only 60 sec or until an infusion was earned (FR1), whichever occurred first. Each animal was trained on only one schedule (either RR or RI). For the RI schedule, the first press on the seeking lever initiated the start of the random interval, and then the first press made following the random interval completed the schedule. Training for the seeking lever began at RR3 or RI10 (criterion of 2 days ≥10 infusions), progressed to RR10 or RI30 (criterion of 2 days ≥10 infusions), and then to the final schedule of RR20 or RI60 (criterion of 5 days ≥10 infusions). The MED-PC program determined the random ratio or interval for a given trial via a probability function (i.e., 0.05 probability per lever press for RR20; 0.0166 probability per second for RI60). Animals were removed from studies if they did not meet the minimum criteria after two weeks at a given stage of training.

### Cocaine outcome devaluation

Once animals were trained on the final seeking-taking schedule (criterion of 5 days ≥ 10 infusions), outcome devaluation was tested across consecutive days in a within-subject manner (devaluation and nondevaluation days, counterbalanced order). As described previously (Jones *et al*, 2022), on the day of outcome devaluation, rats were placed into the operant conditioning chambers and after 5 min, were given experimenter-administered IV cocaine, consisting of 10 μl (to fill the catheter volume) plus a dose that mimicked the estimated brain cocaine concentrations during self-administration; based on the average infusions of the 4 previous self-administration sessions, rats were given 1.0 mg/kg (average of 11-17 infusions during self-administration), 1.5 mg/kg (18-24 infusions), or 2.0 mg/kg (25-31 infusions), in increments of 0.5 mg/kg infusions separated by 20 sec. After a 60-sec waiting period, the seeking lever was available with S+ for 10 min under extinction conditions. On the day of nondevaluation, no infusions were administered but animals spent a similar amount of time in the chamber prior to starting the 10-min extinction test. Devaluation and nondevaluation responding was normalized per rat, such that the number of lever presses on one session was divided by the total lever presses on both sessions (e.g., devaluation lever presses / (devaluation lever presses + nondevaluation lever presses)). Each 10-min devaluation or nondevaluation test was followed by a 5-min period with no levers extended and then the start of a typical cocaine self-administration session. For acclimation purposes, at least 2 days prior to the first devaluation test, rats were given a 10-min extinction session similar to the nondevaluation day. If animals failed to meet a criterion of ≥10 presses during the nondevaluation test session, then both the devaluation and nondevaluation sessions were repeated; if they failed again, then they were removed from analyses.

### Food self-administration

Separate groups of rats were trained to self-administer food pellets (45-mg plain purified pellets, Bio-Serv, Flemington, NJ). Rats were mildly food-restricted for the entire experiment but still gained weight. Rats were fed in the home cage each day >1 h after the operant conditioning session ended and were fed the maximum amount possible that also resulted in all food eaten before the next day’s session (∼50 g for males, ∼20 g for females). Rats underwent the same training as described above for cocaine with a seeking-taking chained schedule of reinforcement (RR20 or RI60), except that a press on the taking lever resulted in delivery of a food pellet paired with tone and light cues. Rats experienced a time out between trials (although only 1 min and the seeking lever was retracted). Sessions were limited to 1 h or 30 rewards, whichever occurred first, so that the total trials per session were comparable to cocaine studies.

### Food outcome devaluation

Once animals were trained on the final seeking-taking schedule (criterion of 5 days with ≥20 rewards), outcome devaluation was tested in a within-subject manner via sensory-specific satiety (devaluation and nondevaluation days, counterbalanced order). Rats were allowed to free-feed on either 45-mg plain purified food pellets (for the devaluation day; the same pellets earned during self-administration) or 15% sucrose solution (for the nondevaluation day) in the home cage for 1 h prior to being placed into the operant conditioning chamber. After a 5-min waiting period, the seeking lever was available for 10 min under extinction conditions, and then rats were returned to the home cage. A normal self-administration session took place the next day, and then the second test (devaluation or nondevaluation, depending on counterbalanced order) took place the following day. Devaluation and nondevaluation responding was normalized per rat, as described above for cocaine outcome devaluation. Rats were acclimated to 15% sucrose by giving it in the home cage overnight ≥ 3 days prior to the first devaluation test.

### Footshock punishment

Once animals were trained on the final seeking-taking schedule (≥ 13 days for cocaine, ≥ 7 for food), and ≥ 4 days after outcome devaluation, they received four consecutive days of punishment sessions. Rats received a minimum of 26 total cocaine self-administration sessions, or 21 total food self-administration sessions, prior to punishment testing. During these sessions, footshock (0.4 mA, 0.3 seconds) was administered on 1/3 trials randomly, after completion of the seeking link and before extension of the taking lever. Rewards were still available on footshock trials. After the 4 days of punishment, rats returned to daily self-administration and were allowed to recover to baseline responding levels (≥ 4 sessions with ≥ 10 rewards) before outcome devaluation testing.

### Shock sensitivity threshold testing

Rats were tested for shock sensitivity before and after all experimentation (with the exception of two male rats in cocaine group). Rats were placed into operant chambers and given a series of footshocks (0.3 s duration, ≥ 10 s inter-shock interval), in an ascending series of intensities from 0.1 to 1.0 mA in 0.1-mA steps. The rats were scored for their first flinch, jump, and vocalization to the footshock, as described by Maren et al (1994). Following vocalization, the ascending series was repeated twice more, and scores across the three test sessions were averaged.

### Data analyses

Animals were removed from all analyses if they failed to meet the criteria for self-administration (described in Methods). Data were analyzed using t-tests or 2-way ANOVAs (with repeated measures when appropriate) as detailed in the Results, with Sidak’s multiple comparisons tests used for post hoc analyses; post hoc results are shown on figures. Statistical results are reported only for effects with significant *p* values (< 0.05). K-means clustering analysis was used to identify and separate sensitive and insensitive groups. Correlation analyses were evaluated via the Pearson correlation coefficient (*r*). Figures show means ± SEM.

## Results

### Cocaine self-administration

Male and female rats were trained on a seeking-taking chained schedule of self-administration for IV cocaine (2 h per day) and then exposed to 4 days of punishment testing. For each rat, the 4 days before punishment were used as a baseline to assess the effects of punishment. Rats that completed ≥65% of baseline trials in the fourth punishment session were considered punishment resistant, whereas rats that completed <65% were considered punishment sensitive. We established this threshold of 65% based on a larger population of male rats exposed to punishment and k-means clustering analysis identifying two clusters with a consistent split at 65% (Fig. S1). A subset of these male rats is included in the following analyses.

Sensitive and resistant rats were significantly different in terms of percent trials completed during punishment, for both males (Fig. 1a; 2-way ANOVA: Group *F*_1,26_ = 26.3, *p* < 0.0001; Day *F*_11,286_ = 34.4, *p* < 0.0001; Group x Day interaction *F*_11,286_ = 9.32, *p* < 0.0001) and females (Fig. 1d; 2-way ANOVA: Group *F*_1,24_ = 24.3, *p* < 0.0001; Day *F*_11,264_ = 24.0, *p* < 0.0001; Group x Day interaction *F*_11,264_ = 7.05, *p* < 0.0001). Sensitive and resistant rats also differed in terms of total trials completed during punishment, but not before punishment, for males (Fig. 1b; 2-way ANOVA: Group *p* = 0.09; Day *F*_11,286_ = 33.1, *p* < 0.0001; Group x Day interaction *F*_11,286_ = 9.63, *p* < 0.0001) and females (Fig. 1e; 2-way ANOVA: Group *F*_1,24_ = 7.70, *p* = 0.01; Day *F*_11,264_ = 22.8, *p* < 0.0001; Group x Day interaction *F*_11,264_ = 6.30, *p* < 0.0001). Males and females were not significantly different from each other in terms of punishment for cocaine, when comparing percent trials completed on the fourth day of punishment for all rats (*t*_52_ = 0.76, *p* = 0.45; male average 57.8% vs. female 52.2%).

**Fig. 1.**
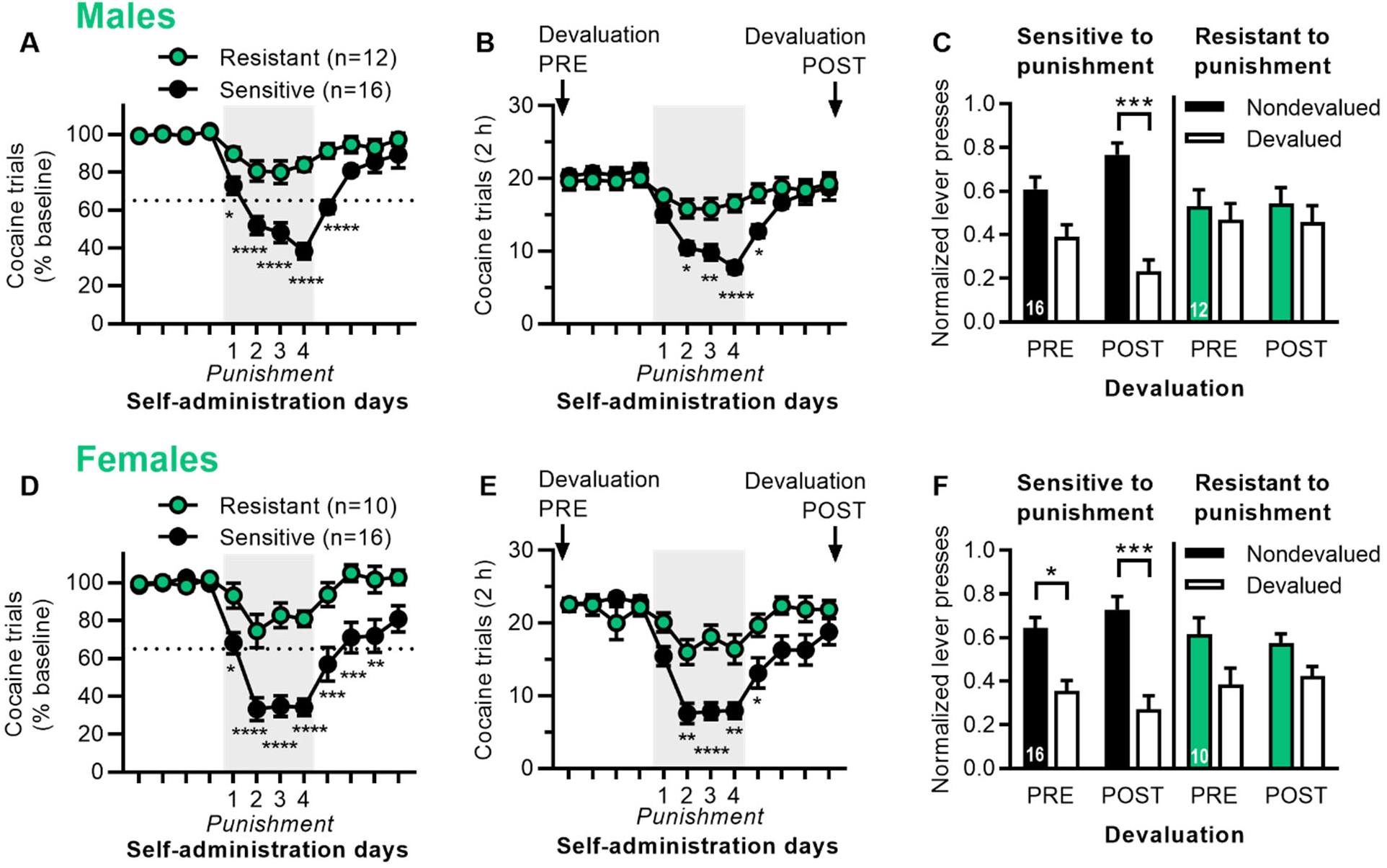
Punishment resistance for cocaine self-administration is associated with inflexible habits. A) Cocaine trials (% baseline) for the 4 days before, during, and after punishment testing for male rats categorized as punishment resistant or sensitive. B) Cocaine trials (total in 2 h) for male rats. Labels indicate when outcome devaluation was conducted pre- and post-punishment. C) Outcome devaluation pre- and post-punishment for male rats that were sensitive or resistant to punishment. Normalized lever presses are shown for nondevalued and devalued sessions. D) Cocaine trials (% baseline) for the 4 days before, during, and after punishment testing for female rats categorized as punishment resistant or sensitive. E) Cocaine trials (total in 2 h) for female rats. F) Outcome devaluation pre- and post-punishment for female rats that were sensitive or resistant to punishment. *p* values < *0.05, **0.01, ***0.001, ****0.0001.

We used outcome devaluation via cocaine satiety to assess whether responding was goal-directed or habitual 4 days pre-punishment and at least 4 days post-punishment (once rats had recovered from punishment), with each rat given devaluation and nondevaluation sessions in a counterbalanced order. In male rats (Fig. 1c), both punishment sensitive and resistant rats were insensitive to outcome devaluation pre-punishment, indicating habitual responding. However, punishment sensitive rats showed increased sensitivity to outcome devaluation post-punishment, indicating enhanced goal-directed control, while punishment resistant rats remained habitual (2-way ANOVA: Devaluation *F*_1,52_ = 12.7, *p* < 0.001; Devaluation x Group interaction *F*_3,52_ = 3.16, *p* = 0.03). In female rats (Fig. 1f), punishment sensitive rats were sensitive to outcome devaluation pre- and post-punishment, while punishment resistant rats were insensitive to outcome devaluation pre- and post-punishment, indicating habitual behavior (2-way ANOVA: Devaluation *F*_1,48_ = 22.5, *p* < 0.0001). Raw data for devaluation testing is shown in Fig. S2.

We then determined whether habitual responding was predictive of punishment resistance. When rats were classified as goal-directed (< 0.4 for normalized devalued responding) or habitual (≥ 0.4) based on pre-punishment outcome devaluation, there was no difference in terms of baseline responding or the fourth day of punishment, for male rats (Fig. 2a; 2-way ANOVA: Session *F*_1,26_ = 58.5, *p* < 0.0001; Session x Strategy interaction *p* = 0.15) or female rats (Fig. 2d; 2-way ANOVA: Session *F*_1,24_ = 77.9, *p* < 0.0001; Session x Strategy interaction *p* = 0.54). Similarly, when classified as goal-directed or habitual based on post-punishment outcome devaluation, there was also no difference in terms of baseline responding or the fourth day of punishment, for male rats (Fig. 2b; 2-way ANOVA: Session *F*_1,26_ = 57.0, *p* < 0.0001; Session x Strategy interaction *p* = 0.13) or female rats (Fig. 2e; 2-way ANOVA: Session *F*_1,24_ = 82.2, *p* < 0.0001; Session x Strategy interaction *p* = 0.16). However, we found that punishment resistance was correlated with habits post-punishment, but not pre-punishment. Specifically, responding on the fourth day of punishment (% baseline trials) correlated with devalued responding during outcome devaluation conducted post-punishment in males (Fig. 2c; *r* = 0.35, *p* = 0.069) and females (Fig. 2f, *r* = 0.43, *p* = 0.029). In contrast, punishment did not correlate with devalued responding conducted pre-punishment in males (*r* = 24, *p* = 0.23) or females (*r* = -0.02, *p* = 0.92). Interestingly, habitual responding was not required for punishment resistance, and some male and female rats that showed resistance (≥ 65% on x-axis) also showed goal-directed responding post-punishment (< 0.4 on y-axis).

**Fig. 2.**
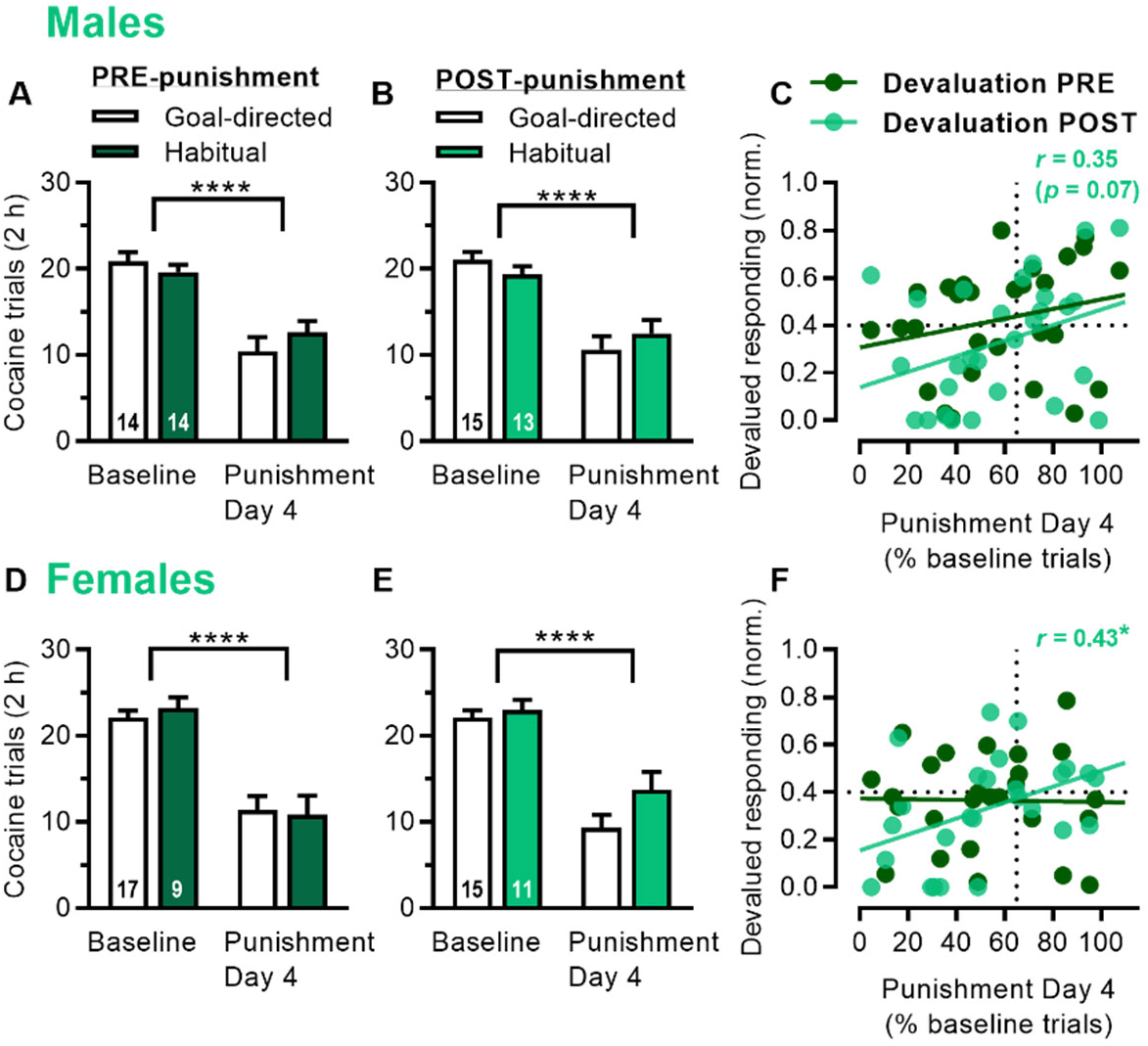
Post-punishment, but not pre-punishment, cocaine habits are associated with punishment resistance. A) Cocaine trials (total in 2 h) during baseline seeking-taking and punishment Day 4 for male rats classified as goal-directed or habitual based on outcome devaluation conducted pre-punishment. B) Cocaine trials for male rats classified as goal-directed or habitual based on outcome devaluation conducted post-punishment. C) Relationship between punishment sensitivity (% baseline trials on Day 4; ≥ 65% threshold considered resistant) and outcome devaluation (normalized devalued responding; ≥ 0.4 threshold considered habitual) for male rats, with devaluation scores pre-punishment (dark green) and post-punishment (light green, *r* = 0.35, *p* = 0.07). D) Cocaine trials (total in 2 h) during baseline seeking-taking and punishment Day 4 for female rats classified as goal-directed or habitual based on outcome devaluation conducted pre-punishment. E) Cocaine trials for female rats classified as goal-directed or habitual based on outcome devaluation conducted post-punishment. F) Relationship between punishment sensitivity and outcome devaluation for female rats, with devaluation scores pre-punishment (dark green) and post-punishment (light green, *r* = 0.43). *p* values < *0.05, ****0.0001.

### Food self-administration

Separate groups of male and female rats were trained on a seeking-taking chained schedule of self-administration for food and then exposed to 4 days of punishment testing. Food self-administration sessions were limited to 1 h or 30 rewards (whichever occurred first), so we used reward rate (pellets per min) to more accurately assess the effects of punishment. For each rat, the 4 days before punishment were used as a baseline. Similar to cocaine, we used a threshold of 65% on the fourth punishment session to identify rats that were resistant to punishment. Sensitive and resistant rats were significantly different in terms of percent baseline during punishment, for both males (Fig. 3a; 2-way ANOVA: Group *F*_1,31_ = 19.9, *p* < 0.0001; Day *F*_11,341_ = 11.9, *p* < 0.0001; Group x Day interaction *F*_11,341_ = 6.68, *p* < 0.0001) and females (Fig. 3d; 2-way ANOVA: Group *F*_1,20_ = 27.9, *p* < 0.0001; Day *F*_11,220_ = 14.0, *p* < 0.0001; Group x Day interaction *F*_11,220_ = 6.53, *p* < 0.0001).

**Fig. 3.**
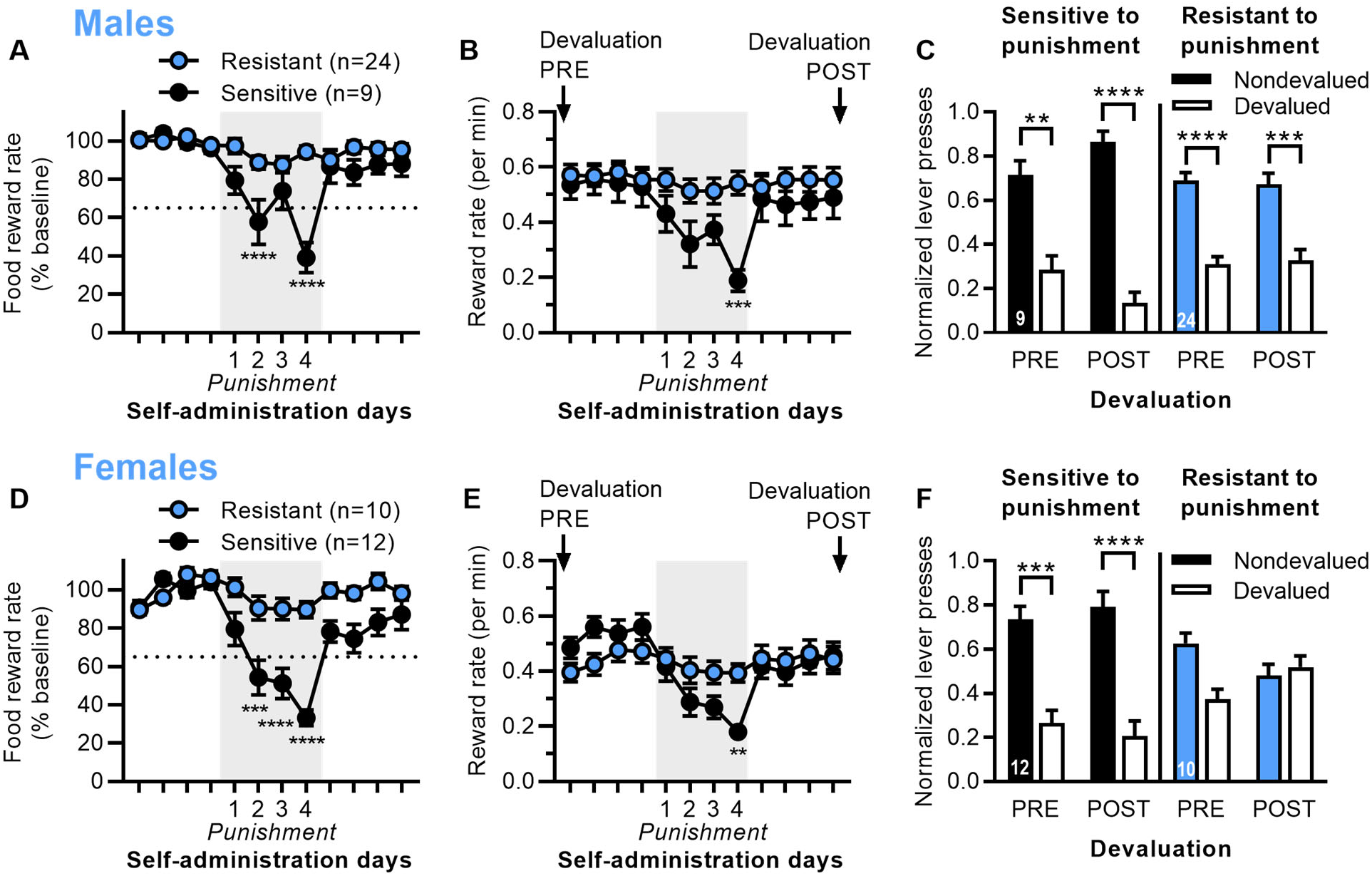
Punishment resistance for food self-administration is associated with inflexible habits, particularly in female rats. A) Reward rate for food (% baseline) for the 4 days before, during, and after punishment testing for male rats categorized as punishment resistant or sensitive. B) Reward rate (pellets/min) for male rats. Labels indicate when outcome devaluation was conducted pre- and post-punishment. C) Outcome devaluation pre- and post-punishment for male rats that were sensitive or resistant to punishment. Normalized lever presses are shown for nondevalued and devalued sessions. D) Reward rate for food (% baseline) for the 4 days before, during, and after punishment testing for female rats categorized as punishment resistant or sensitive. E) Reward rate (pellets/min) for female rats. F) Outcome devaluation pre- and post-punishment for female rats that were sensitive or resistant to punishment. *p* values < **0.01, ***0.001, ****0.0001.

Sensitive and resistant rats also differed in terms of reward rate during punishment, but not before punishment, for males (Fig. 3b; 2-way ANOVA: Group *p* = 0.19; Day *F*_11,341_ = 13.6, *p* < 0.0001; Group x Day interaction *F*_11,341_ = 8.32, *p* < 0.0001) and females (Fig. 3e; 2-way ANOVA: Group *p* = 0.75; Day *F*_11,220_ = 12.9, *p* < 0.0001; Group x Day interaction *F*_11,220_ = 6.65, *p* < 0.0001).

Males were significantly more resistant than females for food, when comparing percent baseline on the fourth day of punishment for all rats (*t*_53_ = 2.4, *p* = 0.020; male average 79.3% vs. female 58.9%). However, there was no significant difference between males and females for baseline reward rate during self-administration (*t*_53_ = 1.5, *p* = 0.13), despite large differences in weight between males and females at the time of punishment testing (*t*_53_ = 19.2, *p* < 0.0001; male average 470 g, female 295 g). We found that males were more resistant for food than cocaine (*t*_59_ = 2.9, *p* = 0.0056), but there was no difference in females for food and cocaine (*t*_46_ = 0.78, *p* = 0.44).

We used outcome devaluation via satiety to assess whether responding was goal-directed or habitual 4 days pre-punishment and at least 4 days post-punishment (once rats had recovered from punishment), with each rat given devaluation sessions (food pellets in home cage) and nondevaluation sessions (15% sucrose in home cage) in a counterbalanced order. In male rats (Fig. 3c), both punishment sensitive and resistant rats were sensitive to outcome devaluation pre- and post-punishment, although sensitive rats showed even greater sensitivity post-punishment, indicating enhanced goal-directed control (2-way ANOVA: Devaluation *F*_1,62_ = 73.6, *p* < 0.001; Devaluation x Group interaction *p* = 0.10). In female rats (Fig. 3f), punishment sensitive rats were sensitive to outcome devaluation pre- and post-punishment, while punishment resistant rats were insensitive to outcome devaluation pre- and post-punishment, indicating habitual behavior (2-way ANOVA: Devaluation *F*_1,40_ = 23.2, *p* < 0.0001; Group *F*_3,40_ = 351, *p* < 0.0001; Devaluation x Group interaction *F*_3,40_ = 5.39, *p* = 0.0033). This mimicked what we observed with cocaine punishment in female rats. Raw data for devaluation testing is shown in Fig. S3.

We determined whether habitual responding was predictive of punishment resistance. When rats were classified as goal-directed (<0.4 for normalized devalued responding) or habitual (≥0.4) based on pre-punishment outcome devaluation, there was no difference in terms of baseline responding on the fourth day of punishment, for male rats (Fig. 4a; 2-way ANOVA: Session *F*_1,31_ = 10.9, *p* = 0.0025; Session x Strategy interaction *p* = 0.60) or female rats (Fig. 4d; 2-way ANOVA: Session *F*_1,20_ = 23.8, *p* <0.0001; Session x Strategy interaction *p* = 0.21). When classified as goal-directed or habitual based on post-punishment outcome devaluation, male rats did not show a significant difference in terms of baseline responding or the fourth day of punishment (Fig. 4b; 2-way ANOVA: Session *F*_1,31_ = 7.46, *p* = 0.010; Session x Strategy interaction *p* = 0.11). However, female rats showed a significant difference for the fourth day of punishment, indicating that the rats classified as habitual post-punishment showed greater punishment resistance (Fig. 4e; 2-way ANOVA: Session *F*_1,20_ = 36.5, *p* < 0.0001; Session x Strategy interaction *F*_1,20_ = 9.64, *p* = 0.0056). Similar to cocaine studies, we found that punishment resistance was correlated with habits post-punishment, but not pre-punishment. Responding on the fourth day of punishment (% baseline reward rate) correlated with devalued responding during outcome devaluation conducted post-punishment in males (Fig. 4c; *r* = 0.39, *p* = 0.027) and females (Fig. 4f, *r* = 0.64, *p* = 0.0014). In contrast, punishment did not correlate with devalued responding conducted pre-punishment in males (*r* = 0.18, *p* = 0.31) or females (*r* = 0.34, *p* = 0.12). Particularly for male rats, habitual responding was not required for punishment resistance, and some male rats that showed resistance (≥ 65% on x-axis) also showed goal-directed responding post-punishment (< 0.4 on y-axis).

**Fig. 4.**
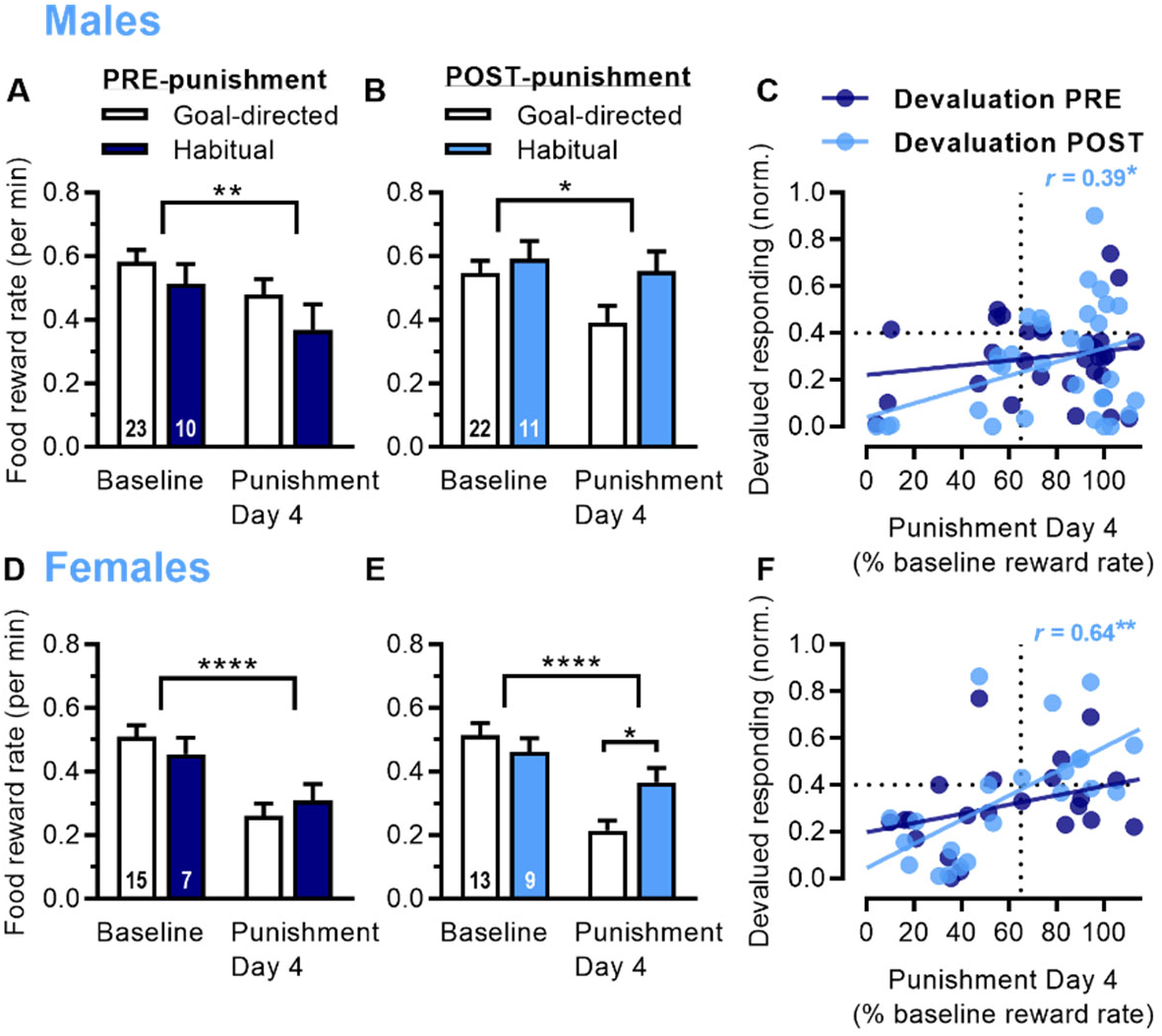
Post-punishment, but not pre-punishment, food habits are associated with punishment resistance. A) Reward rate for food (pellets per min) during baseline seeking-taking and punishment Day 4 for male rats classified as goal-directed or habitual based on outcome devaluation conducted pre-punishment. B) Reward rate for male rats classified as goal-directed or habitual based on outcome devaluation conducted post-punishment. C) Relationship between punishment sensitivity (% baseline reward rate on Day 4; ≥ 65% threshold considered resistant) and outcome devaluation (normalized devalued responding; ≥ 0.4 threshold considered habitual) for male rats, with devaluation scores pre-punishment (dark blue) and post-punishment (light blue, *r* = 0.39). D) Reward rate for food pellets (pellets per min) during baseline seeking-taking and punishment Day 4 for female rats classified as goal-directed or habitual based on outcome devaluation conducted pre-punishment. E) Reward rate for female rats classified as goal-directed or habitual based on outcome devaluation conducted post-punishment. F) Relationship between punishment sensitivity and outcome devaluation for female rats, with devaluation scores pre-punishment (dark blue) and post-punishment (light blue, *r* = 0.64). *p* values < *0.05, **0.01, ****0.0001.

### Correlations with punishment resistance

We ran correlations to determine whether punishment resistance was associated with and/or might be explained by differences in footshock sensitivity or weight. To determine sensitivity to footshock, we conducted footshock threshold testing (threshold for flinch, jump, and vocalization, or FJV) prior to any self-administration training. We found no difference for initial footshock sensitivity between punishment resistant and sensitive groups for cocaine or food in males or females (2-way ANOVAs of FJV vs. group, *p* > 0.05). For cocaine, we found a significant correlation in female rats for vocalization threshold and punishment resistance (*r* = 0.56, *p* < 0.01), but no other correlations in females or males (Fig. 5a, b). For food, we found no correlations between footshock sensitivity and punishment in females or males (Fig. 5 c, d).

**Fig. 5.**
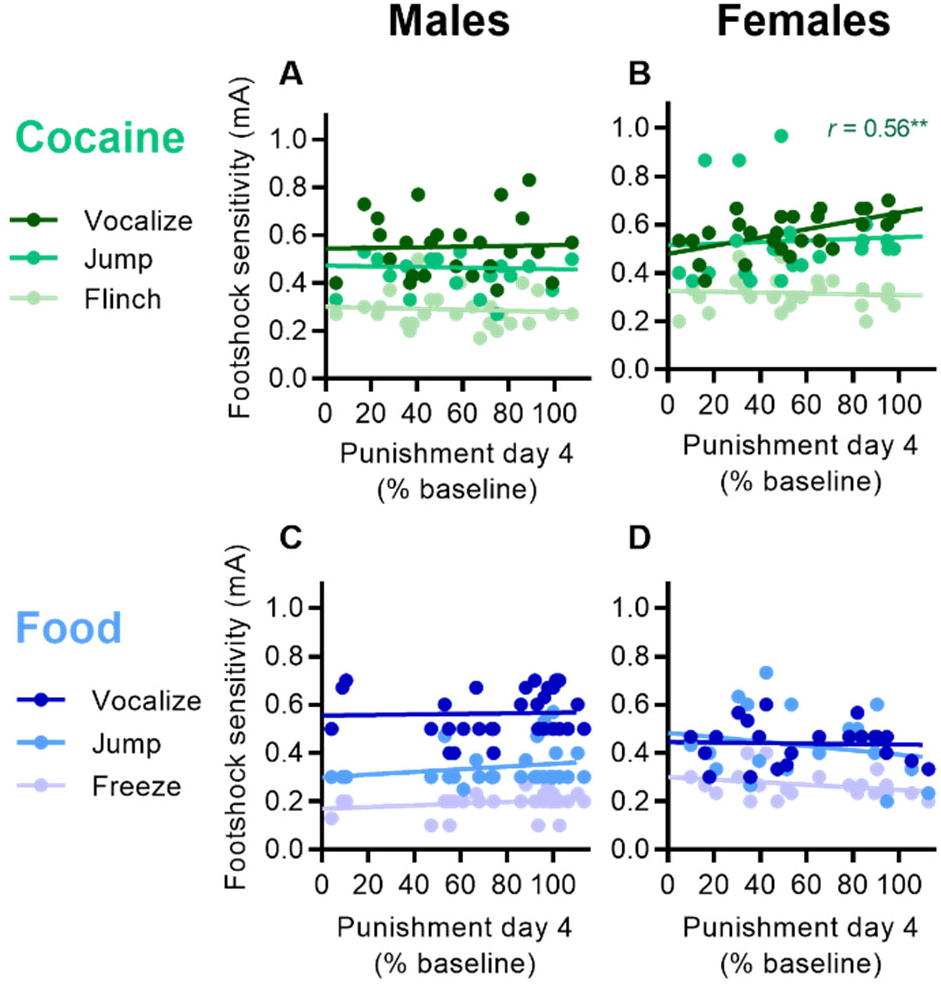
Relationship between punishment resistance and footshock sensitivity (thresholds for flinch, jump, and vocalization). A) Relationship between cocaine punishment sensitivity (% baseline trials) and footshock sensitivity in male rats. B) Relationship between cocaine punishment sensitivity and footshock sensitivity in female rats (vocalization *r* = 0.56). C) Relationship between food punishment sensitivity (% baseline reward rate) and footshock sensitivity in male rats. D) Relationship between food punishment sensitivity and footshock sensitivity in female rats. \*\**p* < 0.01.

We found no significant correlations between punishment sensitivity and animal weight on the first day of punishment; the average weights on the first day of punishment were: cocaine males (540 g), cocaine females (320 g), food males (470 g), and food females (300 g). The average weight gain from the start of self-administration to punishment testing was: cocaine males (200 g), cocaine females (50 g), food males (100 g), and food females (20 g). A comparison of footshock sensitivity testing with the starting weights of the animals revealed a significant correlation only for vocalization in females (*r* = 0.30, *p* = 0.041), but not for flinch or jump in females, or any measure in males, indicating that differences in shock sensitivity were not related to weight.

## Discussion

We found that punishment resistance for cocaine was associated with habitual responding after, but not before, punishment. In other words, habits did not predict punishment resistance, but punishment resistance was related to the continued use of habits. We observed similar results with rats trained to self-administer food, particularly in females. These data indicate that punishment resistance is associated with inflexible habits, whereas punishment sensitivity is associated with increased goal-directed control.

### Punishment resistance and inflexible habits

These findings support the hypothesis that compulsive drug seeking is related to failure to control habits (Belin *et al*, 2013; Everitt and Robbins, 2005, 2016; Everitt, 2014; Ostlund and Balleine, 2008; Smith and Laiks, 2018). While habits themselves are not necessarily maladaptive or permanent, compulsive drug use may be related to a loss of control over habitual seeking, making them inflexible and maladaptive. This idea is corroborated by our finding that habitual reward seeking did not predict punishment resistance. Rather, punishment resistance was related to inflexible habits, whereas punishment sensitivity was related to increased goal-directed control over behavior. Interestingly, habitual behavior was not necessary for punishment resistance and a subset of resistant animals showed goal-directed cocaine seeking. This supports the theory that drug seeking can be a goal-directed choice (Hogarth, 2020; Singer *et al*, 2018).

### Punishment sensitivity and flexible habits

We observed that many rats showed a switch from habitual to goal-directed cocaine seeking when faced with footshock consequences, particularly when sensitive to punishment. Previous studies have also shown animals switching between goal-directed and habitual response strategies. Most commonly, these studies demonstrated a transition from goal-directed to habitual behavior, with the transition occurring gradually over time with progressive training for food (Adams, 1982; Dickinson *et al*, 1995; Holland, 2004; Thrailkill and Bouton, 2015), cocaine (Leong *et al*, 2016; Zapata *et al*, 2010), or alcohol (Corbit *et al*, 2012; Giuliano *et al*, 2021). Similarly, a transition from DMS to DLS control over reward-seeking behavior has been depicted over training (Corbit *et al*, 2012; Murray *et al*, 2012, 2014; Zapata *et al*, 2010). However, several recent studies from Bouton and colleagues demonstrated a transition from habitual to goal-directed responding, instead, and this transition occurred rapidly (Bouton, 2021). The circumstances that caused behaviors to become goal-directed include changes in context, changes in outcome, or unexpected food reinforcers, even when delivered in a different context (Bouton *et al*, 2020; Steinfeld and Bouton, 2020, 2021; Thrailkill and Bouton, 2015; Trask *et al*, 2020). Here, we found that the addition of an aversive outcome enhanced goal-directed responding, but only in animals that were sensitive to footshock punishment. Altogether, these findings indicate that habits generally are not permanent and that the goal-directed system can gain control over behavior.

Typically, habits are somewhat flexible. While habits are insensitive to changes in the value of the outcome, they are sensitive to changes in the outcome. Therefore, habits are not completely inflexible and behavior is typically updated under certain circumstances, including after devaluation when an animal experiences the outcome in the devalued state (Adams, 1982; Balleine *et al*, 2003; Corbit and Balleine, 2003; Ostlund and Balleine, 2008; Yin *et al*, 2005). For example, rats over-trained for food responding showed habitual behavior and insensitivity to outcome devaluation when tested under extinction conditions, but then became sensitive and reduced responding after several minutes of experiencing the food reward actually being delivered (Adams, 1982; Ostlund and Balleine, 2008). Lesions of DMS slowed learning of this effect, indicating that goal-directed processes are typically recruited and that the learning process in the habit system is slower (Ostlund and Balleine, 2008). Ostlund and Balleine (2008) hypothesized that fast changes in performance (e.g., when faced with negative consequences) require a transition to goal-directed control, and that compulsive behavior in addiction may be related to a dominant habit system and difficulty reengaging goal-directed control. The data presented here support this hypothesis, and indicate that the goal-directed and habitual systems function in parallel, such that both DMS and DLS encode the behavior. This explains why post-training inactivation of DMS or DLS leaves behavior intact, but guided by the remaining system (Corbit *et al*, 2012; Giuliano *et al*, 2019; Yin *et al*, 2005, 2006; Zapata *et al*, 2010), and why behavior can rapidly transition to being goal-directed (Bouton, 2021).

We found that the addition of a footshock outcome enhanced goal-directed responding in punishment-sensitive rats. This may seem contradictory to previous work showing that stress biases toward habitual behavior in humans and animals (as reviewed by Schwabe and Wolf, 2011; Smith and Laiks, 2018). However, the impact of stress is dependent on the controllability of the stressor. While inescapable stress has negative long-term effects and recruits the habit system, escapable stress is protective against future insults and recruits the goal-directed system, including DMS and the prelimbic prefrontal cortex (Amat *et al*, 2005, 2006, 2008, 2014; Maier and Watkins, 2010; Maier *et al*, 2006). Previous work has implicated impairments in prelimbic prefrontal function with punishment resistance (Chen *et al*, 2013; Kasanetz *et al*, 2013; Radke *et al*, 2015; Verharen *et al*, 2019). Therefore, it is tempting to speculate that punishment-resistant rats may have a reduced ability to detect control (or contingency) of the footshock (Jean-Richard-Dit-Bressel *et al*, 2019). Further, because prelimbic cortex is necessary for the acquisition of goal-directed responding (Ostlund and Balleine, 2005), reduced prelimbic function might impair the ability to recruit the goal-directed system.

### Reward and sex differences

We observed punishment resistance with both cocaine and food self-administration, and even found that punishment resistance was greater for food than cocaine in male rats. In contrast, previous studies did not observe punishment resistance with food, even in rats with extended sucrose or chow self-administration (Limpens *et al*, 2014; Pelloux *et al*, 2007, 2015; Radke *et al*, 2017; Vanderschuren and Everitt, 2004). In addition, punishment resistance for cocaine was observed with extended, but not limited, exposure to cocaine self-administration (Deroche-Gamonet *et al*, 2004; Jonkman *et al*, 2012a; Pelloux *et al*, 2007; Xue *et al*, 2012), although the pattern of intake is also an important consideration (James *et al*, 2019b; Kawa *et al*, 2016). These discrepancies may be attributed to differences in methods, including footshock intensity, omission of cocaine on footshock trials, schedule of reinforcement, and criteria for resistance.

We found no sex difference in punishment resistance for cocaine, but found increased punishment resistance in males for food, as compared to females. Previous studies have shown female rats to be more sensitive to punishment for cocaine and more sensitive to punishment with risky food rewards (Bender and Torregrossa, 2023; Datta *et al*, 2018a, 2018b; Jacobs and Moghaddam, 2020; Orsini *et al*, 2016). Females also appeared to show greater punishment resistance for cocaine than food (Datta *et al*, 2018a). In contrast, we observed no significant difference for punishment resistance between food and cocaine for females, but found a significant difference between food and cocaine for males. Sex differences in punishment sensitivity for food can likely be traced to sex differences in food motivation and eating. Although we found no sex difference in food reward rate during self-administration, male rats gained more weight than females across the course of the study, and previous work showed that male rats work harder than females to earn food rewards, even despite footshock risk (Jacobs and Moghaddam, 2020; Orsini *et al*, 2016). We found that punishment-resistant rats were insensitive to outcome devaluation pre- and post-punishment in most groups we studied (males and females with cocaine, and females with food). However, male rats with punishment resistance for food were sensitive to outcome devaluation pre- and post-punishment, indicating goal-directed responding. Therefore, punishment resistance for food in male rats was not necessarily related to inflexible habits, and in some animals, seemed to be more related to goal-directed actions.

### Mechanisms of punishment resistance

The current data support the theory that punishment resistance is related to loss of control over habits (Belin *et al*, 2013; Brown *et al*, 2022; Everitt and Robbins, 2005, 2016; Everitt, 2014; Ostlund and Balleine, 2008; Smith and Laiks, 2018). However, it is important to note that support for this theory is not mutually exclusive with other theories of addiction (see commentary by Epstein, 2020). There are multiple factors that contribute to drug seeking and addiction, and the factors may differ across individuals (i.e. individual differences) or have a compound influence within an individual. Several factors have been hypothesized or considered for a possible influence on punishment resistance, including habitual behavior, goal-directed behavior, high value of cocaine, low value for footshock, and reduced contingency learning (Everitt and Robbins, 2016; Field *et al*, 2020; Hogarth, 2020; Jean-Richard-Dit-Bressel *et al*, 2018; Lüscher *et al*, 2020; Smith and Laiks, 2018).

Although we did not find that habits predicted punishment resistance for cocaine or food, previous work showed that punishment-resistant alcohol seeking was greater in rats with DLS-dependent alcohol seeking (Giuliano *et al*, 2018). In addition, punished responding for cocaine was reduced by DLS inactivation (Jonkman *et al*, 2012b), but not by inhibition of DMS direct pathway (Yager *et al*, 2019), further implicating DLS in punishment resistance. In contrast, support for the theory that addiction is related to excess goal-directed motivation comes from work indicating that habits are not prerequisite for punishment resistance (Singer *et al*, 2018), and that resistance was associated with strengthened activity in a pathway between orbitofrontal cortex and DMS (Hu *et al*, 2019; Pascoli *et al*, 2018). In addition, punished responding for food in mice was associated with reward-related dopamine signals in DMS and not DLS, and responding was reduced by DMS manipulations (Seiler *et al*, 2022).

Punishment resistance could be related to increased motivation for the reward or decreased sensitivity to the aversive consequence. In support of the former, punishment resistance was associated with higher break point on a progressive ratio schedule of reinforcement, as well as lower demand elasticity (i.e., high motivation) using behavioral economic measures (Bentzley *et al*, 2014; Deroche-Gamonet *et al*, 2004; James *et al*, 2019a; Kasanetz *et al*, 2010). However, other work showed no association between punishment resistance and break point for cocaine or sucrose rewards, even though higher doses of cocaine drove greater resistance (Datta *et al*, 2018a, 2018b). However, there is little support for punishment resistance being driven by decreased sensitivity to aversive consequences. Punishment-resistant alcohol seeking was not related to differences in footshock-induced fear (Giuliano *et al*, 2018). Further, punishment-resistant cocaine seeking was not correlated with punishment-resistant sucrose seeking, indicating that punishment resistance cannot simply be attributed to individual differences in footshock sensitivity (Datta *et al*, 2018a, 2018b). Further, we found that punishment resistance could not be explained by decreased sensitivity to footshock. Finally, studies using a conditioned punishment task for food rewards found little evidence that punishment resistance is related to reward dominance or aversion insensitivity, and punishment resistance in rats and humans seemed most causally related to a lack of learning the punishment contingency and understanding the relationship between actions and aversive outcomes (Jean-Richard-Dit-Bressel *et al*, 2019, 2023).

### Conclusions and future directions

We found that punishment resistance for cocaine was associated with inflexible habits, whereas punishment sensitivity was associated with exerting goal-directed control. We did not find that habitual responding predicted punishment resistance. However, future studies with extended training on cocaine self-admiration might reveal that habits become even more inflexible and predictive of punishment resistance. Future work might also explore further explore the hypothesis that punishment resistance is related to impaired contingency detection.

## Acknowledgments

The authors thank the many undergraduate researchers that assisted with conducting behavioral studies. This work was supported by National Institutes of Health grant R01 DA046457 (RJS) and start-up funds provided by Texas A&M University.

