## Supplementary Figures for "Punishment resistance for cocaine is associated with inflexible habits in rats"

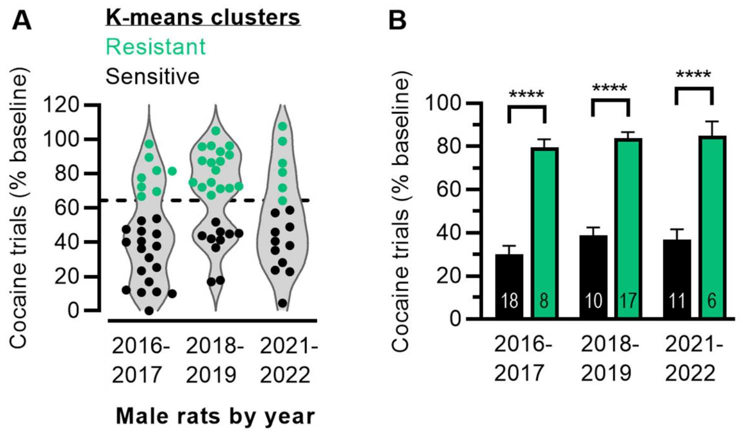


**Fig. S1 | Establishing a threshold for punishment resistance.** A) Violin plots show male rats that received footshock punishment of cocaine self-administration grouped together by year they were ordered. Individual data points represent cocaine trials (% baseline) on the fourth day of punishment for each rat. Within each group, separate K-means clustering analysis identified two clusters (represented by green and black dots), with a consistent threshold of 65% splitting the clusters. The cluster above this threshold is labelled punishment resistant, and the cluster below this threshold is labelled punishment sensitive. B) Mean (± SEM) cocaine trials (% baseline on fourth day of punishment) for sensitive and resistant groups separated by year (2-way ANOVA: Group $F_{1,64} = 171$, $p < 0.0001$; post hoc ****$p < 0.0001$).

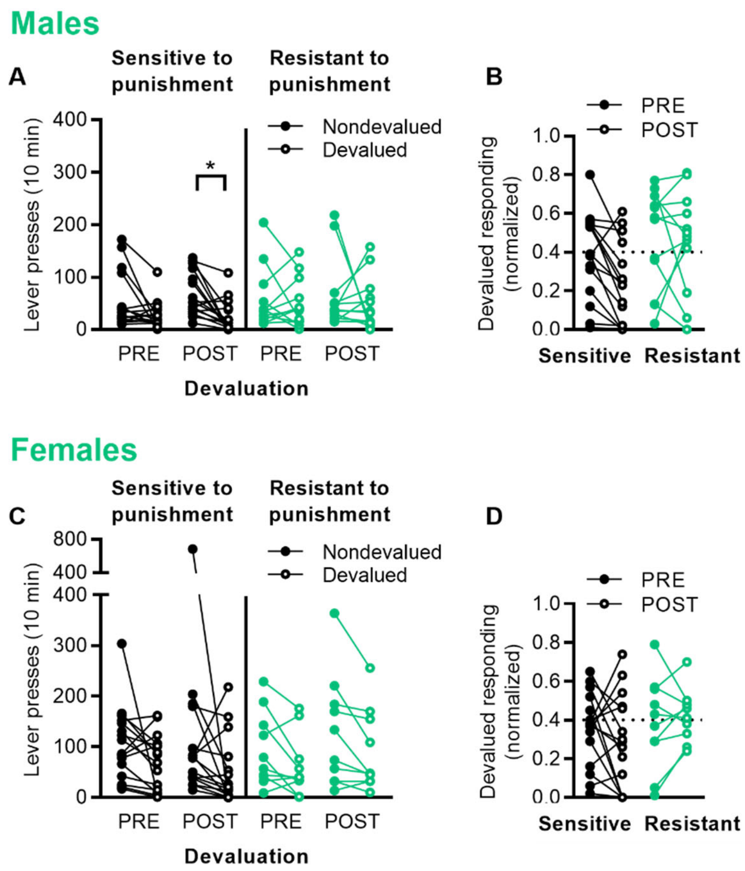


**Fig. S2 | Supplemental data for outcome devaluation in cocaine self-administration.** A) Raw data for outcome devaluation pre- and post-punishment for male rats that were sensitive or resistant to punishment (normalized data is shown in Fig. 1c). Lever presses are shown for nondevalued and devalued sessions with connecting lines for each rat (2-way ANOVA: Devaluation $F_{1,52} = 7.45$, $p = 0.0086$; Devaluation x Group interaction $p = 0.43$; $^*p < 0.05$). B) Comparison of pre- and post-punishment devalued responding scores for individual male rats sensitive or resistant to punishment. Below 0.4 threshold is considered goal-directed; above habitual. C) Raw data for outcome devaluation pre- and post-punishment for female rats that were sensitive or resistant to punishment (normalized data is shown in Fig. 1f). Lever presses are shown for nondevalued and devalued sessions with connecting lines for each rat (2-way ANOVA: Devaluation $F_{1,48} = 7.57$, $p = 0.0084$; Devaluation x Group interaction $p = 0.80$). D). Comparison of pre- and post-punishment devalued responding scores for individual female rats sensitive or resistant to punishment.

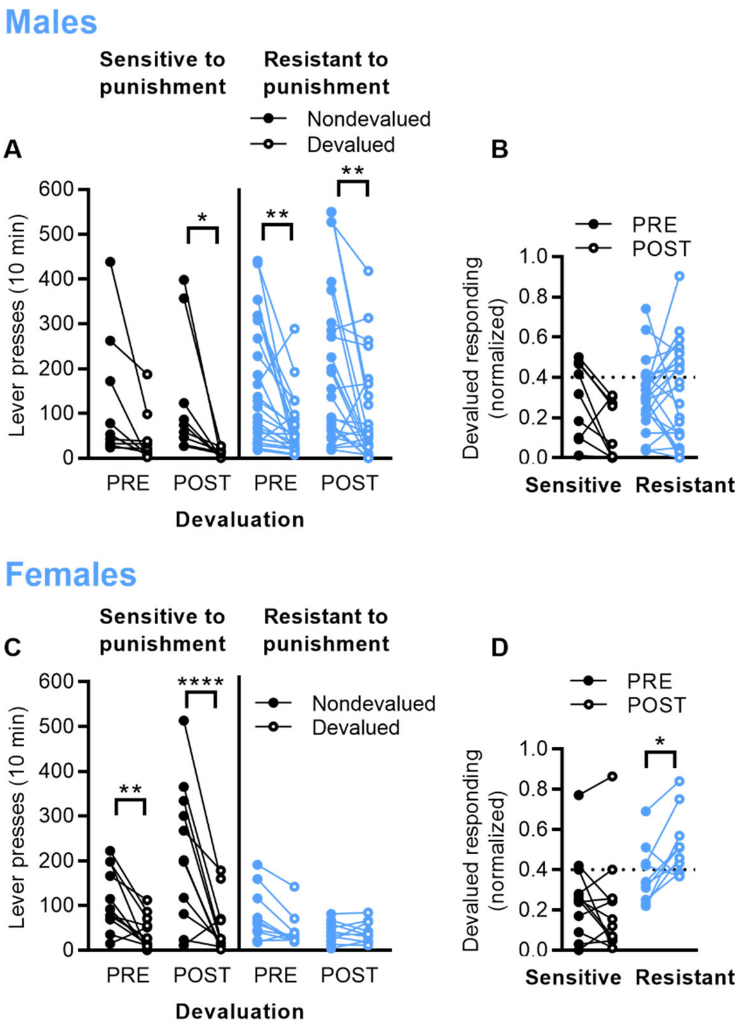


**Fig. S3 | Supplemental data for outcome devaluation in food self-administration.** A) Raw data for outcome devaluation pre- and post-punishment for male rats that were sensitive or resistant to punishment (normalized data is shown in Fig. 3c). Lever presses are shown for nondevalued and devalued sessions with connecting lines for each rat (2-way ANOVA: Devaluation $F_{1,62} = 27.6$, $p < 0.0001$; Devaluation x Group interaction $p = 0.94$). B) Comparison of pre- and post-punishment devalued responding scores for individual male rats sensitive or resistant to punishment. Below 0.4 threshold is considered goal-directed; above habitual. C) Raw data for outcome devaluation pre- and post-punishment for female rats that were sensitive or resistant to punishment (normalized data is shown in Fig. 3f). Lever presses are shown for nondevalued and devalued sessions with connecting lines for each rat (2-way ANOVA: Devaluation $F_{1,40} = 34.6$, $p < 0.0001$; Group $F_{3,40} = 5.08$, $p = 0.0045$; Devaluation x Group interaction $F_{3,40} = 9.90$, $p < 0.0001$). D) Comparison of pre- and post-punishment devalued responding scores for individual female rats sensitive or resistant to punishment (2-way ANOVA: Group $F_{1,20} = 7.63$, $p = 0.012$; Group x Pre/post interaction $F_{1,20} = 9.32$, $p = 0.0063$). $p$ values < *0.05, **0.01, ****0.0001.
